# The Epigenetic Factor PHF13 Governs Trophoblast Stemness and Differentiation

**DOI:** 10.64898/2026.03.14.711150

**Authors:** Sheng Liu, Lei Liu, Jiayu Meng, Elena Sadovsky, Keyi Huang, Heather Sorenson, Tianjiao Chu, Yoel Sadovsky, Yingshi Ouyang

## Abstract

Differentiation of trophoblast stem (TS) cells or progenitor cytotrophoblasts (CTBs) into multinucleated syncytiotrophoblasts (STBs) is essential for placental development. Disruption of this process contributes to major obstetrical syndromes, including fetal growth restriction and preeclampsia, and Trisomy 21. However, the chromatin mechanisms governing trophoblast stemness and differentiation remain inadequately defined. Here we identify the chromatin-associated factor PHF13, uncovered through a high-throughput microRNA target screen, as a key regulator of trophoblast cell fate. *PHF13* knockout TS cells exhibited defects that ultimately resulted in loss of cell viability, whereas *PHF13* knockdown promoted expression of fusion-associated genes, including *ERVFRD-1* and human chorionic gonadotropin (*hCG*). Consistently, *PHF13* depletion in BeWo trophoblast cells increased *hCG* expression and secretion while reducing expression of canonical stemness-associated transcription factors *ELF5* and *TEAD4*. Integrated genomic analyses further revealed that PHF13 target genes comprise a gene regulatory network that maintains trophoblast stemness and restrains differentiation. Notably, the pluripotency-associated transcription factor THAP11 partially co-occupies genomic sites with PHF13. Together, these findings establish PHF13 as a previously unrecognized chromatin regulator of trophoblast stemness and differentiation, providing mechanistic insight into pathways critical for placental development and function.

**Highlights:**

- PHF13 preserves trophoblast stem cell identity
- PHF13 restrains syncytiotrophoblast differentiation
- PHF13 controls trophoblast chromatin programs with THAP11

## INTRODUCTION

The human placenta is vital for intrauterine fetal development and growth, mediating maternal-fetal exchange of gases, nutrients, metabolites, hormones, and extracellular vesicles. Villous trophoblasts, which constitute the forefront fetal tissue that is directly bathed in maternal blood, includes progenitor mononuclear cytotrophoblast (CTB) and differentiated, fused multinuclear syncytiotrophoblasts (STBs). The STB is critical for regulating the maternal-fetal exchange and thus, pregnancy health. Pathobiological processes that emanate from perturbed villous trophoblast differentiation and fusion have been implicated in common pregnancy syndromes^1, 2, 3^, such as subtypes of fetal growth restriction, preeclampsia, and chromosomal abnormalities such as Trisomy 21^4^, which collectively represent major causes of pregnancy-related morbidity and mortality worldwide.

The balance of trophoblast self-renewal and differentiation is pivotal for the development of human trophoblast, a process broadly modulated by two distinct gene categories: (1) transcriptional regulators that establish and maintain trophoblast identity and stemness, and (2) genes that promote trophoblast differentiation and fusion. Stemness-associated transcription factors include Caudal type homeobox 2 (*CDX2*), GATA binding protein 2/3 (*GATA2/3*), E74 like E26 transformation-specific transcription factor 5 (*ELF5*), transcriptional enhancer factor-1 domain transcription factor 1 and 4 (*TEAD1/4*) and their cofactor *YAP1*, transcription factor AP-2 alpha/gamma (*TFAP2A/C*), and tumor protein p63 (*TP63*)^5, 6, 7, 8, 9, 10^. In contrast, trophoblast differentiation is characterized by activation of fusion-promoting genes, including transcription factor glial cells missing transcription factor 1 (*GCM1*), human chorionic gonadotropin (*hCG*), and endogenous retroviral (ERV) envelope proteins such as *ERVW-1* (*aka syncytin-1*) and *ERVFRD-1 (aka syncytin-2)*, as well as their cognate receptors *SLC1A5* (aka *ASCT2*) and *MFSD2a*^11, 12, 13, 14, 15, 16^. In the trophoblastic stem/progenitor state, stemness genes are robustly expressed, whereas fusion-promoting genes are repressed. Conversely, differentiation is accompanied by transcriptional repression of trophoblast stemness programs, with augmented expression of genes promoting fusion and endocrine maturation.

The balance between trophoblast stemness, differentiation, and syncytium formation is also influenced by epigenomic modifications. Recent advances in the studies of histone methylation and acetylation have deepened our understanding of how these histone codes contribute to transcriptional regulation in the healthy and diseased placenta. Specifically, increased expression of histone 3 lysine 4 trimethylation (H3K4me3) and histone 3 lysine 27 acetylation (H3K27ac) are associated with preeclampsia^17, 18^. Further, H3K4me3 and other histone codes are implicated in trophoblast differentiation, exhibiting highly complex occupancy changes in a variety of genes across gestation^18, 19, 20^. While these observations suggest that genomic regions demarcated by histone marks may account for trophoblastic gene expression in CTB *vs* STB, the key epigenomic factors that mediate their effects in distinct trophoblastic cell states remain incompletely defined.

The development of human trophoblast stem (TS) cells, derived from multiple sources^21, 22, 23, 24, 25^, provides an invaluable experimental system for investigating the molecular pathways that regulate the balance between trophoblast stemness and differentiation. Importantly, TS cells exhibit three key trophoblastic features^26, 27, 28^: (1) trophoblast progenitor properties, including the expression of a pan-trophoblast marker cytokeratin 7 and trophoblastic stemness genes; (2) differentiation bi-potential to villous or to HLA-G-expressing extravillous trophoblasts *in vivo* and *in vitro*; and (3) the expression of villous trophoblast-specific the chromosome 19 microRNA cluster. In this study we focused on the role of Plant Homeodomain Finger 13 (PHF13) in human trophoblast biology. PHF13 was identified through our high-throughput RNA-based screening approach^29, 30^, and has been characterized as a chromatin-associated protein implicated in reproductive development^31^. Here we demonstrate that PHF13 promotes a TS-CTB stemness state and suppresses villous trophoblastic differentiation. Together, these findings uncover PHF13 as a previously underappreciated chromatin-associated regulator that links epigenomic control to transcriptional networks governing trophoblast stemness and differentiation.

## RESULTS

### PHF13 expression in the human placenta

In the pursuit of factors that mediate the regulatory functions of trophoblast-specific miRNAs encoded by the human chromosome 19 microRNA cluster (C19MC), we deployed the Crosslinking, Ligation and Sequencing of Hybrids (CLASH) technology^29^ to identify RNA targets for C19MC miRNA. We found that *PHF13* is one of the top transcripts targeted by C19MC miR-517a-3p^30^. PHF13 has been implicated in host cell response to viral infections^32, 33^, and in sustaining spermatogonia stem cells^34^. Mechanistically, through binding to histone modification H3K4me3, PHF13 controls gene expression in mouse embryonic stem cells^31^. Based on our data and the observation that H3K4me3 exhibits profound occupancy alterations in mononucleated CTB *vs* multinucleated STB^18, 19, 20^, we sought to interrogate a potential role of PHF13 in human trophoblasts. We first assessed the expression of *PHF13* transcripts in term placentas of uncomplicated pregnancies. Employing RNAscope technology, we found that *PHF13* was expressed in the trophoblast layer (Suppl. Fig. 1A-C). We also detected *PHF13* mRNA in non-trophoblastic villous cells, which is aligned with its ubiquitous expression pattern, as documented in the Human Protein Atlas. We further located PHF13 protein expression mainly to the syndecan1-demarcated trophoblast layer of term placentas from uncomplicated pregnancies (Suppl. Fig. 1D-E).

### PHF13 impact on trophoblastic transcriptomes

As primary human trophoblasts can be maintained only several days in culture, they are not accessible to CRISPR-based technology. We therefore utilized TS cells in studies of trophoblast differentiation^26^. Note that TS cells were rigorously defined, exhibiting *in vivo* differentiation into two main trophoblast lineages^21^. To interrogate PHF13 function in human trophoblasts, we generated *PHF13* knockout (ko) TS cells using CRISPR as we recently detailed^30^. T7 endonuclease assays confirmed the ko clones, producing the expected genomic fragments.

Compared to wild type (wt) cells, *PHF13* ko cells exhibited cell death in two independent clonal cells (Suppl. Fig. 2). Unable to expand these cells, we could not validate PHF13 protein levels in the ko clones.

To circumvent the cell lethality caused by inactivation of *PHF13* in TS-CTB cells, we generated *PHF13* ko in the commonly used trophoblast line BeWo, and confirmed depletion of PHF13 protein in these ko BeWo cells (Fig. 1A). Unlike *PHF13* ko TS cells, *PHF13* ko BeWo cells remain viable, enabling functional analysis of PHF13 in trophoblast differentiation. Using hCG production and secretion as established markers of STB differentiation, we found that *PHF13* ko cells exhibited markedly increased *hCG* mRNA levels and hCG protein release (Fig. 1B), indicating activation of a differentiation program. Consistent with this finding, differentiation-promoting genes, including transcription factor *GCM1*, its downstream target *ERVFRD-1*, and ERVFRD-1’s cognate receptor MFSD2a, were upregulated in *PHF13* ko cells, whereas the expression of stemness genes, including *ELF5, TEAD4, and TP63*, was reduced in PHF13 ko cells (Fig. 1C). These results suggest that loss of PHF13 promotes trophoblast differentiation.

**Fig. 1:**
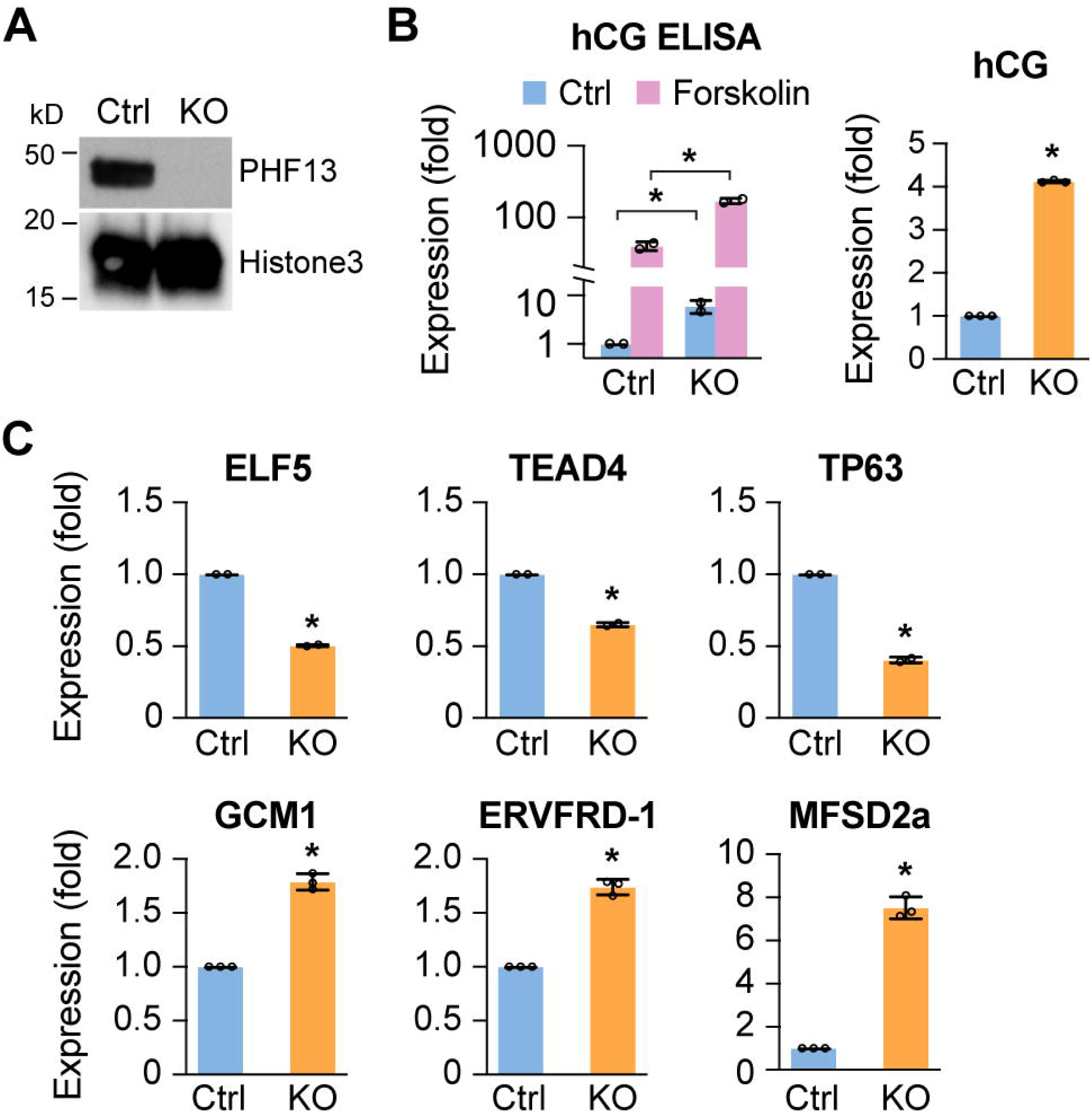
Loss of PHF13 potentiates trophoblast differentiation. **A.** PHF13 ko in BeWo cells, validated by western blot. Histone H3 was used as a loading control. **B.** PHF13 ko increases medium hCG release (ELISA) and expression (qPCR). BeWo cells were exposed to 50 μM forskolin for 72 h to induce differentiation. **C.** The effects of PHF13 ko on stemness genes including *ELF5*, *TEAD4*, and *TP63*, and fusion-promoting genes, including *GCM1*, *ERVFRD-1* and its receptor *MFSD2a*. N=3, *p < 0.01, t-test.

To define the global transcriptional impact of PHF13 deficiency, we performed RNAseq and identified 2,456 differentially expressed genes in *PHF13* ko *vs* wt BeWo cells. Consistent with the increased hCG production, *PHF13* ko led to upregulation of multiple STB-dominant genes, including *CYP19A1*, *syndecan 1, and ERVFRD-1*, with no effect on *ERVW-1* expression (Suppl. Table 1). Together, these results suggest that PHF13 is necessary for the expression of canonical stemness genes while repressing activation of differentiation and fusion-promoting genes in BeWo cells. Therefore, loss of PHF13 releases a transcriptional brake on trophoblast differentiation, permitting induction of STB-related gene networks.

Because complete inactivation of *PHF13* resulted in loss of TS-CTB cells, we generated two shRNA knockdown lines (*PHF13* kd1 and kd2 hereafter) and analyzed their transcriptome by RNAseq. As expected, the expression of *PHF13* mRNA were significantly decreased (log_2_fold change (FC): -1.35, Suppl. Table 2). Compared to control cells, *PHF13* kd1 led to altered expression of 1072 genes, including 549 downregulated genes and 523 upregulated genes, whereas *PHF13* kd2 resulted in 3490 differentially expressed (DE) genes, including 1654 downregulated genes and 1836 upregulated genes (Suppl. Table 2). Using the Pearson’s linear correlation and Spearman’s rank correlation, we confirmed the correlation between the transcriptional landscape of *PHF13* kd1 and *PHF13* kd2 TS-CTB cells, when compared to controls (Suppl. Fig. 3A). Comparing expression changes between the two independent PHF13 kd TS-CTB cells, we found concordant changes in 621 genes, including 321 downregulated genes and 300 upregulated genes (Suppl. Fig. 3B). Further gene ontology analysis revealed the prominent enriched terms, ranked in descending order of enrichment scores (Suppl. Fig. 3C).

Notably, pathways related to epithelial branching morphogenesis and actin filament assembly, which are associated with trophoblast differentiation and syncytial remodeling, were significantly enriched. These included gene ontology terms: cell-substrate junction, focal adhesion, cell-to-cell junction, ruffle defined as actin-rich membrane protrusions, and actin binding (Suppl. Fig. 3C).

To test whether PHF13 kd shifts the trophoblastic transcriptomes toward the differentiation state, we applied gene set enrichment analysis (GSEA) using curated sets of 20 representative stemness-related genes (*CDX2, ELF5, TET1, TET2, TFAP2A/2C, KRT7, TEAD1, TEAD3, TEAD4, PEG10, WNT6, SP6, MSX2, GATA2/3, HAND1, YAP1, SMARCA1, and SMARCA4/BRG1) and 20* STB-related genes (*CGA, CGB3, CGB5, CGB8, CRH, CYP11A1, CYP19A1, ERVW-1, ERVFRD-1, GCM1, HMOX1, HSD11B2, LGALS13, LGALS14, LGALS16, OVOL1, PGF, PSG3, PSG11, and TBX3).* GSEA revealed selective enrichment of the STB-dominant gene set in PHF13 kd cells, whereas trophoblastic stemness-related genes showed no significant enrichment (Fig. 2A). These results suggest that PHF13 suppresses the differentiation program in TS cells. Although core stemness transcription factors, including *GATA2/3, ELF5, TEAD1/4, TFAP2A/C* were not affected by *PHF13* kd in TS cells, we found that *WNT6* and *SOX15,* highly expressed in trophoblast progenitor cells^35, 36^, were consistently downregulated in both *PHF13* kd cells (Fig. 2B). This suggests that PHF13 selectively maintains a subset of stemness-associated genes in TS cells. Intriguingly, CDX2, a transcription factor governing early trophectoderm specification, was upregulated at both mRNA and protein levels in *PHF13* kd cells (log_2_FC: 1.63, Suppl. Fig. 4), suggesting that PHF13 normally restrains a trophectoderm-like gene program in TS cells.

**Fig. 2:**
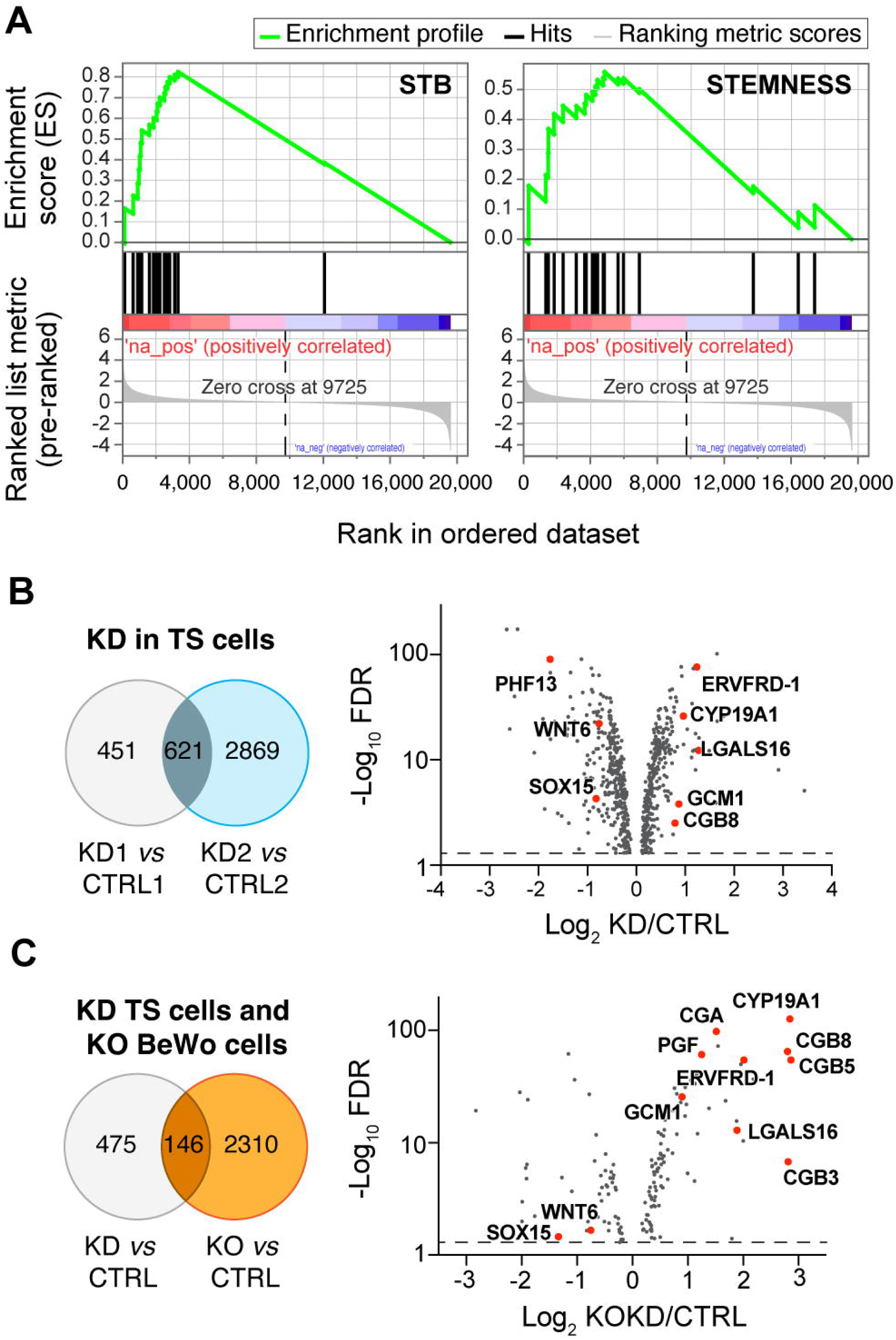
The transcriptomic alterations upon PHF13 knockdown in TS cells. **A.** Gene set enrichment analysis of canonical trophoblastic progenitor genes (shown as STEMNESS, normalized enrichment score: 1.44; FDR-adjusted value: 0.07) and syncytiotrophoblast-dominant genes (shown as STB, normalized enrichment score: 2.11; FDR-adjusted value: 1 x 10^-5^) in TS cells deficient for *PHF13*. Key gene names and results are detailed in the text. **B.** A volcano plot showing the 621 differentially expressed mRNAs shared in both PHF13 kd lines, when compared to controls. The x-axis represents the log_2_ fold change in expression (kd/ctrl), whereas the y-axis shows the minus log_10_ of the FDR-adjusted p value. The horizontal dash line marks the signficance threshold (FDR-adjusted p <0.05; –log_10_ > 1.30). Red points highlight downregulated genes, including *PHF13, WNT6,* and *SOX15*, and upregulated genes that are largely expressed in differentiated trophoblasts, including *ERVFRD-1, CYP19A1, LGALS16, GCM1,* and *CGB8.* **C.** A volcano plot showing the 146 differentially expressed genes shared between *PHF13* kd TS cells and ko BeWo cells. The x-axis represents log_2_ fold change in expression (kokd/ctrl), whereas the y-axis shows the minus log_10_ of the FDR-adjusted p value. The horizontal dash line marks the signficance threshold (FDR-adjusted p <0.05; –log_10_ > 1.30). Red points indicate significantly altered genes, which include downregulated stemness-associated genes (*WNT6* and *SOX15*), and upregulated differentiated trophoblast markers (e.g. *ERVFRD-1, CYP19A1, LGALS16, GCM1, CGA, CGB5, CGB3, CGB8, and PGF*).

Among canonical STB-related transcripts, we found upregulation of *CGA, CGB3, CGB5, CGB8, CYP19A1*, *ERVFRD-1*, *GCM1*, *HMOX1*, *LGALS16*, and *PGF* in both *PHF13* kd cells. Additionally, transcription factor TBX3, which promotes hCG expression^37^, was increased in *PHF13* kd cells (Suppl. Table 2). Notably, corticotropin releasing hormone (CRH), which augments human trophoblast fusion and is exclusively expressed in STB^38, 39, 40^, was also elevated (Log_2_FC: 2.16, Suppl. Table 2) in *PHF13* kd TS cells. Together, our findings uncover a new role of PHF13 in repressing the expression of a myriad of differentiation-promoting, STB-relevant genes in TS cells. Further, intersecting our RNAseq data from *PHF13* kd TS cells and ko BeWo cells, we found 146 overlapping genes (Fig. 2C and Suppl. Table 3). Specifically, stemness genes such as *WNT6* and *SOX15* were downregulated, whereas genes that enhance trophoblast differentiation, such as *GCM1* and *ERVFRD-1*, or genes that are largely expressed in STB, including *PGF, hCG, CYP19A1,* and *LGALS16*, were upregulated (Fig. 2C).

### Target binding by trophoblastic PHF13

Having established the transcriptional consequences of PHF13 deficiency, we next interrogated whether PHF13 directs trophoblastic gene expression by binding to chromatin regulatory regions of its targets^31^. Deploying native chromatin immunoprecipitation via Cleavage Under Targets & Release Using Nuclease and sequencing (CUT&RUNseq), we analyzed PHF13-bound regions in two distinct cell states, TS-CTB and differentiated TS-STB. As a positive control, H3K4me3, which is known to co-bind the nucleosome with PHF13^31^, occupied 12,540 loci in TS-CTB and 9,243 loci in TS-STB. We found that 1,075 and 2,342 genes were targeted by PHF13 in TS-CTB and TS-STB, respectively (Suppl. Table 4).

We first focused on canonical trophoblast stemness genes, including *ELF5, TEAD4, and TP63,* that are downregulated in *PHF13* KO BeWo cells *vs* controls, as well as in TS-STB *vs* TS-CTB cells. We hypothesized that PHF13 occupancy at the genomic loci of *ELF5, TEAD4, and TP63* was required for their transcription in TS cells, and that a loss of PHF13 would reduce their expression. When compared to a control immunoglobin (IgG) immunoprecipitation, PHF13 exhibited significant binding to the genomic loci of *ELF5, TEAD4,* and *TP63* in TS-CTB, with no binding difference in TS-STB (Fig. 3A-C). Unlike *TEAD4*, *TEAD1* genomic loci harbored PHF13 peaks in both mononucleated and multinucleated cell states compared to their respective controls (Fig. 3D). We also found that *WNT6* contained PHF13 peaks in TS-CTB, but not in TS-STB (Fig. 3E), which correlated with its downregulation in *PHF13*-deficient TS-CTB. Consistent with these data, H3K4me3 showed major peaks in the *ELF5, TEAD4, TP63,* and *WNT6* genomic loci in TS-CTB, but not in TS-STB. Together, these results suggest that PHF13 is needed for the expression of *ELF5, TP63, TEAD4,* and *WNT6* in the stemness state.

**Fig. 3:**
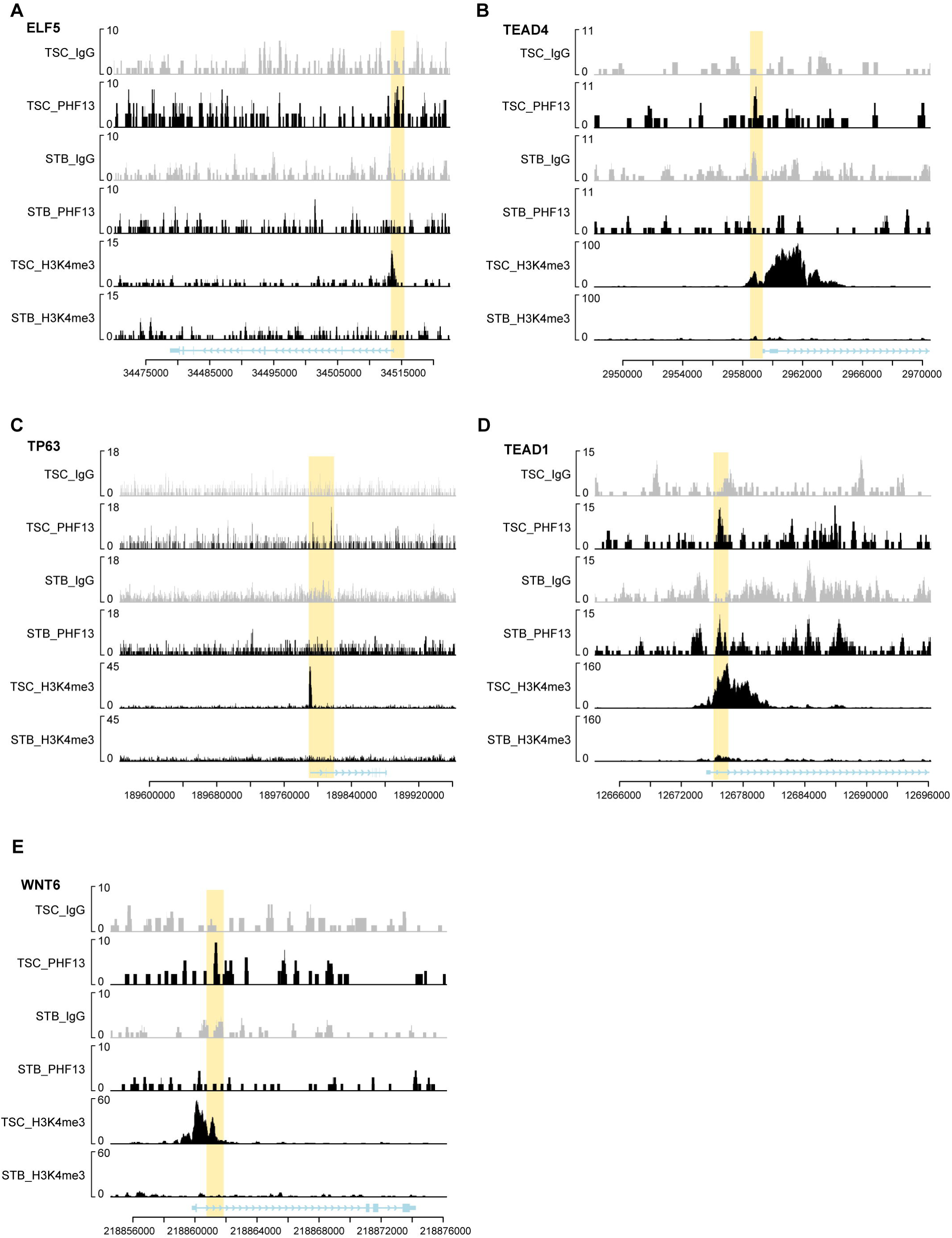
PHF13 binding tracks at trophoblastic stemness genes and TS-associated genes in TS-CTB and TS-STB cells. PHF13-bound regions at trophoblastic stemness genes, including *ELF5* (A), *TEAD4* (B), *TP63* (C), and *TEAD1* (D), and TS-associated genes such as *WNT6* (E). Yellow shading denotes signficant PHF13 peaks (A-C, E) in TS-CTB, but not in TS-STB when compared to corresponding IgG controls. In contrast, *TEAD1* (D) harbored PHF13 peaks in both TS-CTB and TS-STB. As a positive control, H3K4me3 showed binding in all the five TS-related genes in TS-CTB rather than TS-STB (A-E). The x-axis represents genomic corordinates with exonic and intronic structure, and arrows indicate the direction of transcription. The y-axis represents normalized peak intensity.

Having shown that PHF13 activates stemness genes through chromatin binding, we next examined PHF13 occupancy of canonical STB genes that are induced in TS-STB. In TS-STB, PHF13 peaks were detected at *CGA* and *CGB8* (Fig. 4A-B), but not at *CYP19A1*, *GCM1*, *ERVW-1*, or *ERVFRD-1*, when compared to controls (not shown). We then tested whether PHF13 binding in TS-STB preferentially marks genes that are upregulated in TS-STB vs TS-CTB. Of the 2,342 PHF13 targets identified in TS-STB, 288 overlapped with genes upregulated in TS-STB (Suppl. Table 5), representing a 1.34-fold enrichment with p value 1.99 x 10^8^. No enrichment was observed among downregulated genes in TS-STB. Further, Analysis of the NIH Database for Annotation, Visualization and Integrated Discovery (DAVID) of the 288 upregulated PHF13 targets in TS-STB highlighted oxygen-related response pathways, including response to decreased oxygen levels (GO:0036293), response to hypoxia (GO:0001666), and response to oxygen levels (GO:0070482). These targets include N-myc down-regulated gene 1 (NDRG1), which we previously showed is upregulated by hypoxia and protects trophoblasts against hypoxic injury and fetal growth restriction^41, 42, 43, 44^, as well as the hypoxia-responsive non-coding RNA *metastasis-associated lung adenocarcinoma transcript 1 (MALAT1)*^45,46^.

**Fig. 4:**
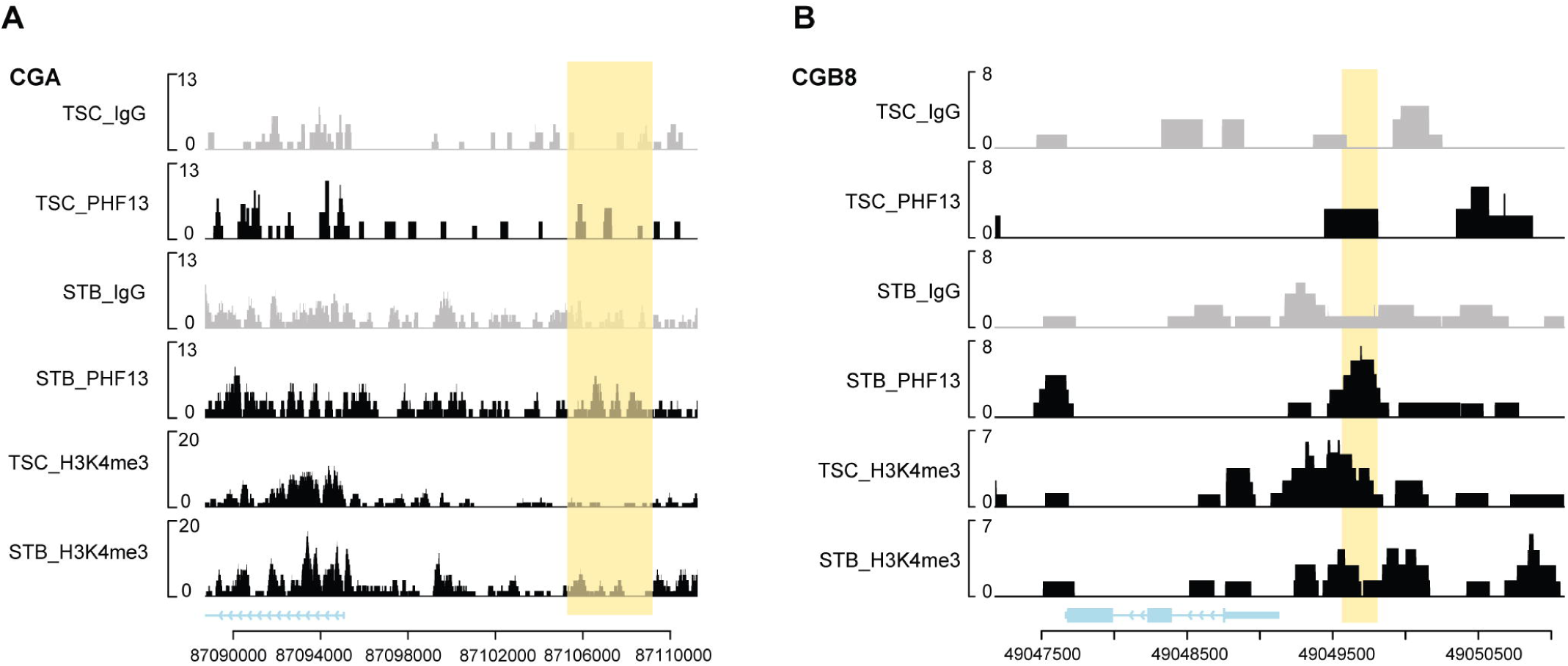
PHF13 binding tracks at differentiation marker hCG subunit loci in TS-CTB and TS-STB cells. Yellow shading denotes PHF13 peaks at the *CGA* (A) and *CGB8* (B) regions in TS-STB but not in TS-CTB, relative to IgG controls. The x-axis represents genomic corordinates with exonic and intronic structure, and arrows indicate transcription direction. The y-axis represents normalized peak intensity.

### Integrated analysis of PHF13 CUT&RUNseq and PHF13 knockdown in TS cells

To determine whether PHF13 binding in TS-CTB marks genes that change expression in *PHF13* kd TS-CTB, we integrated the analysis of PHF13 CUT&RUNseq with *PHF13* kd RNAseq in these cells. Using the over-representation algorithms^47^, we identified 37 downregulated and 48 upregulated genes, whose loci contained PHF13 peaks (Fig. 5A-B). DAVID analysis of the 37 downregulated PHF13 targets showed no enrichment of KEGG or Reactome pathways, but indicated several differentiation-related GO terms, including “cell junction” (GO: 0030054), “cell surface receptor signaling pathway” (GO:0007166), “actin cytoskeleton organization” (GO:0030036), and “response to a cytokine” (GO:0034097) (Suppl. Table 6A). Notably, *IFITM3*, a known inhibitor of trophoblast fusion^48, 49^, was among the 37 downregulated PHF13 targets. We hypothesized that PHF13 promoted *IFITM3* expression by binding its promoter in TS cells. Supporting this hypothesis, we found that PHF13 occupied the *IFITM3* locus in TS-CTB (Fig. 5C), and IFITM3 expression decreased upon PHF13 knockdown (Fig. 5D-E).

**Fig. 5:**
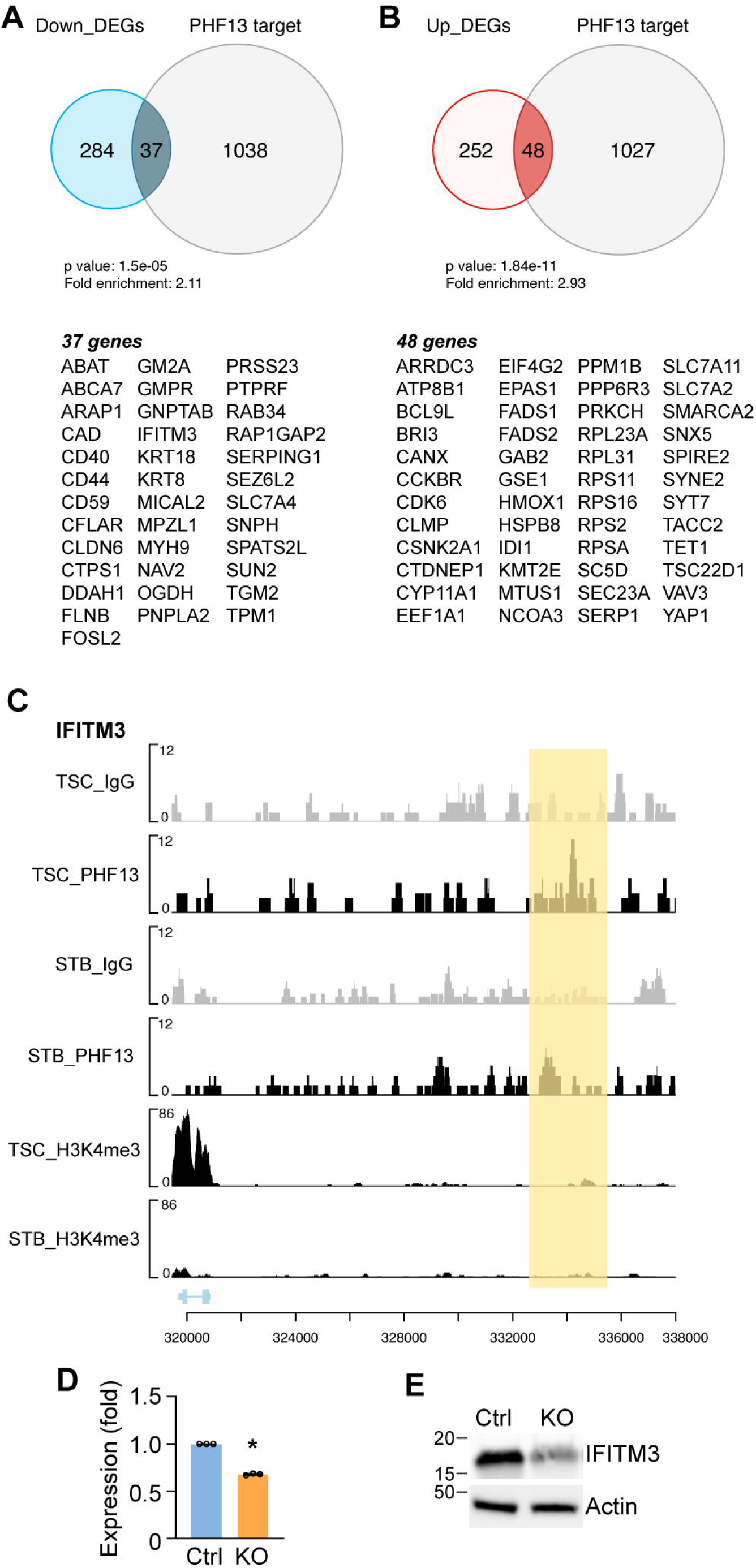
Enrichment analysis of PHF13 target genes and gene expression changes in TS-CTB cells. **A-B.** 37 downregulated genes and 48 upregulated genes in PHF13-deficient cells are direct targets of PHF13. Note that their enrichment is signficant with 2.11-fold and 2.93-fold enrichment, respectively. **C.** PHF13 binds genomic regions of *IFITM3*, an inhibitor of trophoblast fusion, in TS-CTB but not in TS-STB. Yellow shading indicates significant PHF13 peaks relative to IgG control. The x-axis represents genomic corordinates with exonic and intronic structure, and arrows indicate transcription direction. The y-axis represents normalized peak intensity. **D-E.** PHF13 promotes the expression of IFITM3 transcripts (D) and protein (E) in *PHF13* ko BeWo cells. Actin was used as a loading control. N=3, * p < 0.05, t-test.

Analysis of the 48 PHF13-target genes upregulated upon PHF13 loss showed enrichment for stress-response pathways (Suppl. Table 6B). HMOX1, a stress-inducible cytoprotective enzyme important for antioxidative defense and hypoxic adaptation in trophoblasts^50^, was increased in *PHF13* kd TS cells (log_2_FC: 0.93, Suppl. Table 2). Consistently, Human Protein Atlas single-cell RNAseq show lower HMOX1 expression in progenitor CTB than in differentiated STB, supporting a repressive role of PHF13 in controlling *HMOX1* expression in TS-CTB. Among the 27 upregulated genes annotated under “positive regulation of cellular process” (GO:0048522), cholecystokinin B receptor (*CCKBR*), highly expressed in human trophectoderm (TE) prior to implantation^51^, but absent from the inner cell mass^52^, was also a PHF13 target. Notably, *CCKBR* expression was elevated in *PHF13* kd TS-CTB (log_2_FC: 1.16), indicating that PHF13 restrains CCKBR expression in TS-CTB. Consistent with CCKBR’s TE-restricted expression, this suggests that PHF13 helps maintain the TE-TS cell hierarchy by preventing premature activation of a TE program in TS cells. Because the human trophectoderm lineage shows increased ribosome biogenesis and protein synthesis programs compared to embryonic stem cells^52^, we next examined translation-related genes and identified seven PHF13 targets involved in ribosome-related processes, including cap-dependent translation initiation and elongation (Suppl. Table 6B).

Together, these results support a model in which PHF13 maintains a trophoblast stem-like state through chromatin binding at stemness-related gene loci and at the fusion inhibitor IFITM3, while repression of differentiation-related genes such as *GCM1* and *ERVFRD-1* occurs indirectly. Further analysis of PHF13-regulated loci revealed additional state-specific targets, including the opioid agonist precursor gene *PENK*. PHF13 occupied the *PENK* locus and sustained its expression in TS-CTB, whereas in differentiated TS-STB, PENK expression was maintained through PHF13-independent mechanisms (Suppl. Fig. 5 and Suppl. Note 1).

### THAP11 co-occupies with genomic loci of PHF13 target genes

This context-dependent nature of PHF13-mediated gene regulation raised the possibility that PHF13 may need additional cofactors including transcription factors to modulate target gene expression in human trophoblasts. To identify putative cofactors associated with PHF13-bound chromatin, we performed motif enrichment analysis of PHF13 CUT&RUNseq data. This analysis uncovered significant enrichment of the consensus motif for thanatos-associated protein domain containing 11 (THAP11) (Fig. 6A). Because THAP11 is a transcription factor expressed in bovine TS-like cells^53^ and is also required for pluripotency in mouse embryonic stem (mES) cells^54^, we examined its role in human trophoblast. THAP11 protein levels were 50% lower in TS-STB than in TS-CTB (Fig. 6B), suggesting an inverse association with trophoblast differentiation. To assess whether THAP11 influences trophoblastic gene programs, we silenced *THAP11* expression, using Cas13d-mediated RNA knockdown. Given the rapid growth arrest and loss of viability reported for *Thap11* (also called *Ronin*) knockout mES cells^54^, we used knockdown rather than knockout. We confirmed that THAP11 protein was reduced by approximately 90% in kd cells (Fig. 6C). RNAseq revealed that THAP11 reduction had a stronger effect on gene expression during differentiation than under stemness conditions. In kd TS cells, 401 genes were downregulated and 180 upregulated (Suppl. Table 7A), whereas under differentiation conditions, 1,756 were downregulated and 1,916 upregulated (Suppl. Table 7B). Interestingly, differentiation-related genes such as *ERVFRD-1* (log_2_FC: 0.49) and *GCM1* (log_2_FC: 0.33) were upregulated in THAP11-deficient TS-STB cells (Suppl. Table 7B), although their expression in TS-CTB cells were not influenced by THAP11. Together, THAP11 loss alters baseline expression and reshapes the transcriptional response to differentiation.

**Fig. 6:**
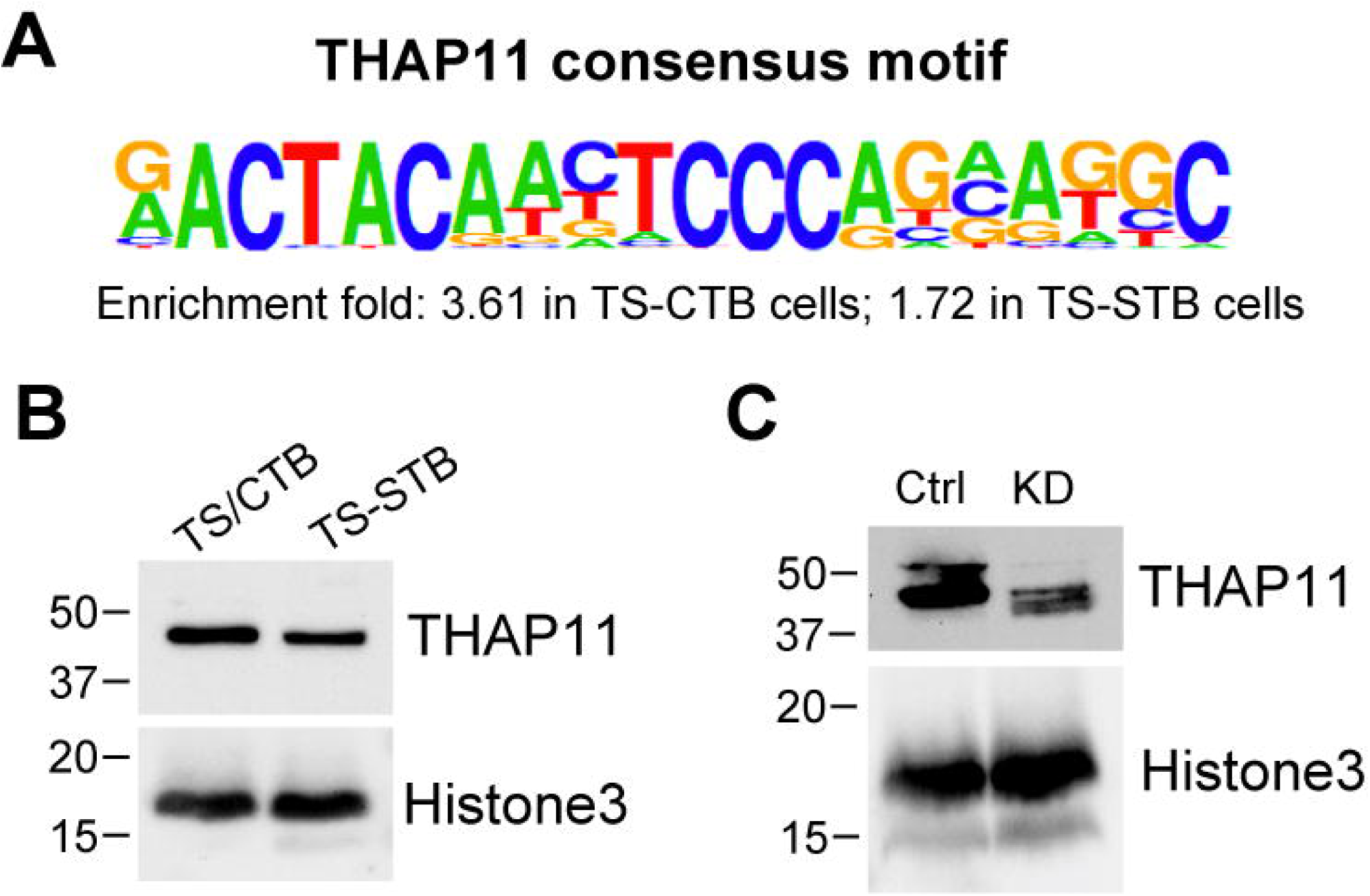
THAP11 is a cofactor for PHF13 to modulate gene expression. **A.** The genomic sequence bound by PHF13 is also a consensus motif recognized by transcription factor THAP11. **B.** THAP11 protein expression is reduced in TS-STB when compared to TS-CTB cells. **C.** A confirmation of THAP11 protein kd. Histone H3 was used as a loading control. N=3, * p < 0.05, t-test.

We next asked which THAP11-responsive genes are targeted by PHF13. Under stemness conditions, 36/401 (9.0%) downregulated genes and 10/180 (5.6%) upregulated genes were PHF13-occupied (Suppl. Table 8A). In contrast, during differentiation, this overlap increased to 234/1,756 (13.3%) downregulated genes and 303/1,916 (15.8%) upregulated genes (Suppl. Table 8B). Function analysis indicated limited pathway-level enrichment. In TS cells, we only found 22/36 (61.1%) PHF13-targeted genes that were downregulated in THAP11 kd cells mapped to the GO cellular component “cell periphery” (GO:0071944); most encode plasma membrane proteins, with the remaining two genes *LAMA1* and *VIT* annotated as extracellular matrix glycoproteins (Suppl. Table 8A). Under differentiation conditions, PHF13-targeted genes that were downregulated in THAP11 kd cells showed enrichment for the biological process “cytoskeleton organization” (GO:0007010; 41/234 genes), and were also enriched for cellular component, including “cell periphery” (GO:0071944; 98/234 genes) and “endomembrane system” (GO:0012505; 92/234 genes in Suppl. Table 8B). In contrast, PHF13-targeted 303 upregulated genes in differentiation medium were enriched for cellular component “cytosol” (GO:0005829; 140/303 genes, in Suppl. Table 8B). Together, these results suggest that the loss of THAP11 perturbs PHF13-linked transcriptional effects, particularly those related to cellular architecture (membrane/endomembrane compartments and cytoskeleton organization) during trophoblast differentiation.

## DISCUSSION

We identified a previously unrecognized regulatory role for PHF13 in human trophoblasts. PHF13 support the stem-like state in TS-CTB cells while restraining differentiation. When the expression level of PHF13 is reduced, cells lose stemness and upregulate genes associated with differentiation, including genes enriched in multinucleated STBs. PHF13 target genes in TS-CTB cells comprise gene regulatory networks that lead to promotion of trophoblastic stemness and inhibition of differentiation. Consistent with this role, PHF13 regulates canonical stemness factors, including *ELF5, TEAD4,* and *TP63*, and supports the expression of additional stemness-associated genes such as *WNT6* and *SOX15*. In TS-CTB cells, PHF13 also targets *IFITM3*, an inhibitor of trophoblast differentiation and fusion. Accordingly, IFITM3 expression decreases upon PHF13 loss.

Not all differentiation-associated genes appear to be directly bound or regulated by PHF13. For instance, *ERVFRD-1* is upregulated in PHF13 kd TS-CTB cells, yet we did not detect PHF13 occupancy at the *ERVFRD-1* locus, suggesting that indirect repression of *ERVFRD-1* by PHF13. This prompted us to surmise that PHF13 acts through distinct binding partners in different cell states. Supporting this possibility, *ERVFRD-1* is strongly silenced by the repressive H3K9me3 mark catalyzed by SETDB1 methyltransferase^55, 56^, and PHF13 has been reported to interact with SETDB1 and promote H3K9me3^57^.

We further examined the regulation of transcription factor CDX2 in the context of PHF13 function. *Cdx2* is essential for mouse trophectoderm specification and embryo implantation, but TS cells, derived from placental villi after 6 gestational weeks^22^, show minimal *CDX2* expression, consistent with low CDX2 abundance at this stage^58, 59^. In contrast, TS cells^21^ express CDX2, enabling us to assess PHF13-dependent regulation. Several lines of evidence suggest that PHF13 indirectly represses *CDX2*. We found that PHF13-demarcated peaks were not enriched at the *CDX2* promoter regions. Additionally, although murine Elf5 is necessary and sufficient for *Cdx2* transcription via binding to the *Cdx2* promoter^60, 61^, upregulation of human *CDX2* in PHF13-deficient cells is unlikely driven by ELF5, which is decreased in PHF13 knockout cells. Moreover, we found that histone acetylase KAT8, reported to promote *CDX2* expression via acetylation of histone H4 at lysine 16 in TS cells^62^, was not altered in *PHF13* kd TS cells or *PHF13* ko BeWo cells. Instead, because *CDX2* expression increases with hypomethylation^63^, and because TET1/2, an enzyme that catalyzes 5-hydroxmethylcytosine formation and is vital for TS cell maintenance^64, 65^, are upregulated in *PHF13* kd TS cells, we suggest that increased TET1/2 may contribute to upregulation of *CDX2* in human TS cells.

Our transcriptomic analyses demonstrate that PHF13 exerts context-dependent regulatory effects on gene expression, with *PHF13* inactivation also leading to upregulation of PHF13-responsive genes. The repressive function of PHF13 in trophoblasts aligns with the evidence that PHF13 promotes local repressive chromatin by facilitating a histone repressive mark H3K9me3^57^, and by enhancing the prominent polycomb repressive complex 2 (PRC2)-dependent H3K27me3 occupancy at selected loci^31^. Therefore, by occupying promoters and regulatory elements, PHF13 can simultaneously sustains stemness maintenance and constrain differentiation-associated transcription. The state-specific regulation of the opioid precursor PENK further exemplifies this context-dependence. Although PENK’s role in trophoblast stemness *vs* differentiation remains unknown, a recent study indicates downregulation of *PENK* upon osteoblastic differentiation^66^. Further, as PENK is vital for generating agonists that activate OPRD1, whose expression is influenced by differentiation^26^, these data suggest that PHF13 might regulate PENK-dependent pathways in TS-CTB cells.

Finally, we identified a subset of genes that are likely coregulated by THAP11 and PHF13. These genes may contribute to remodeling of cellular architecture during trophoblast differentiation. Supporting this notion, loss of THAP11 disrupts expression of genes linked to cell periphery integrity and endosomal organization. Although the precise mechanisms remain to be elucidated, our results suggest that THAP11, along with PHF13, is involved in maintaining gene expression programs that underpin physical and organelle architecture as mononucleated trophoblasts transition through differentiation. Whether PHF13 and THAP11 interact directly, and whether PHF13 chromatin occupancy is required to recruit THAP11 to target loci, remain to be determined.

*Limitations of our study:* This study primarily examined PHF13 function in TS cells and the trophoblast cell line BeWo. Validation of these findings in additional TS cell models^22, 23, 24, 25^, as well as in primary trophoblasts^26, 67^, will be important to confirm their common physiological relevance. Although many genes with altered expression in *THAP11* kd cells are also PHF13 targets, the molecular mechanisms by which THAP11 functions as a PHF13 cofactor to regulate gene expression remain to be defined. For instance, direct biochemical demonstration of PHF13-THAP11 interaction and cooperative chromatin regulation will be an important focus of future studies.

Another important question is whether PHF13 expression or function is altered in pregnancies complicated by fetal growth restriction or preeclampsia, in conditions where trophoblast differentiation is impaired. Given THAP11’s established role in stem cell maintenance, the PHF13-THAP11 chromatin axis identified here thus warrants further investigation to determine how this pathway governs trophoblastic stemness and differentiation balance, and whether disruption of this pathway contributes to placental insufficiency.

## METHODS

### Term placental biopsies with uncomplicated pregnancies

Participants delivered at the Magee-Women’s Hospital (MWH), Pittsburgh, Pennsylvania, and provided an informed consent for placental biopsy collection (protocol #19120076 & 19100240), which was obtained from the Steve N. Caritis Magee Obstetric Maternal and Infant Database and Biobank at Magee-Women’s Research Institute. Placental biopsies (5 mm^3^) were collected immediately after delivery by the obstetrical specimen procurement unit at MWH. The biopsies were obtained from a region that is midway between the cord insertion and the placental margin and between the chorionic and basal plates, as we previously detailed^68^.

### Immunofluorescence of placenta formalin-fixed paraffin-embedded (FFPE) tissues

FFPE sections (5 μm) were mounted on adhesive coverslips, heated at 60°C for 60 min, deparaffinized (xylene, graded ethanol series from 100% to 50%), and rehydrated. Antigen retrieval was performed in basic buffer (10 mM Tris, 1 mM EDTA, pH 8.5) in 95°C water bath for 20 min, followed by cooling at room temperature. Coverslips were then mounted on CellScape Tissue Chips (Canopy Biosciences, #PRSM-CHP-TISSUE-010), washed with PBS and blocked with 1% BSA for 30 min. Imaging and staining were performed on the CellScape instrument (Canopy Biosciences), with autofluorescence captured in each channel and subtracted during analysis. The tissue-containing chips were incubated overnight at 4°C with rabbit anti-PHF13 (Sino Biological, #202944-T08, 1:200 dilution), washed, and then incubated for 30 min at room temperature with PE donkey anti-rabbit (Biolegend, #406421, 1:200), PE anti-human CD138, #356504, 1:300), and Hoechst 33342 (Thermo Fisher, #H3570, 1:1,200 dilution) using microfluidics. Images were acquired and analyzed with the CellScape scanner.

### Cell and culture

All cell lines were maintained in a humidified incubator at 37°C, 20% O_2_ and 5% CO_2_. Human induced trophoblast stem (TS) cells were cultured on collagen-coated dishes in TS medium as previously described^21, 26^. Briefly, collagen IV powder (Sigma, #C5533-5MG) was reconstituted in 5 ml of 0.5% penicillin–streptomycin phosphate-buffered saline (Thermo Fisher, #14190144). The solution was mixed gently using a pipette, incubated overnight at 4°C to ensure complete reconstitution, aliquoted, and stored at -20°C as 1 mg/ml stock solution. Culture plates were precoated with 5 μg/ml collagen IV for at least 1 h at 37°C prior to TS cell seeding.

TS cell medium consisted of basal medium DMEM/F-12, GlutaMAX (Thermo Fisher, #10565042) supplemented with 0.3% bovine serum albumin (BSA; Sigma, #A9576-50ML), 0.2% fetal bovine serum (FBS; Thermo Fisher, #16141079), 1% ITS-X supplement (Thermo Fisher, #51500056), 0.1 mM 2-mercaptoethanol (Thermo Fisher, #31350010), 0.5% penicillin–streptomycin (Thermo Fisher, #15140122), 1.5 μg/ml L-ascorbic acid (Sigma, #A4544-25G), 50 ng/ml epidermal growth factor (EGF; PeproTech, AF-100-15), 1 μM SB431542 (Selleck Chemicals, #S1067), 0.5 μM A83-01 (Sigma, #SML0788-5MG), 0.8 mM valproic acid (Sigma, #P4543-10G), 2 μM CHIR99021 (Miltenyi Biotec, #130-104-172), 5 μM Y-27632 (Selleck Chemicals, #S1049). TS cells were fed with the new TS medium every 3 days and passaged using TrypLE express enzyme, no phenol red (Thermo Fisher, #12604013) and replated onto collagen-coated dishes as described here.

To induce differentiation into syncytiotrophoblasts, TS cells were seeded in a 12-well plate coated with 2.5 μg/ml collagen IV at a density of 300,000 cells per well, and cultured in the differentiation medium, including DMEM/F-12, GlutaMAX (Thermo Fisher, #10565042), 0.3% BSA, 1% ITS-X supplement, 0.1 mM 2-mercaptoethanol, 0.5% penicillin–streptomycin, 4% knockout serum replacement (Thermo Fisher, #10828028), 2 μM forskolin (Selleck Chemicals, #S2449) and 2.5 μM Y-27632.

Human choriocarcinoma BeWo cell lines (ATCC, #CCL-98) were cultured in F-12K Kaighn’s modified medium (ATCC, #30-2004), supplemented with 10% FBS (ATCC, #30-2020) and 1% penicillin–streptomycin and were fed with the new F-12K complete medium every 3 days and passaged using TrypLE express enzyme, phenol red (Thermo Fisher, #12605010).

### Lentivirus production

HEK293T/17 cells (ATCC, #CRL-11268) were seeded at 9 x 10⁶ per 10-cm culture dishes containing DMEM, high glucose, GlutaMAX (Thermo Fisher, #10569044) supplemented with 10% FBS (Thermo Fisher, #16141079) and incubated overnight at 37°C in a 20% O_2_ and 5% CO_2_ incubator. On the following day, the medium was replaced with 10 ml fresh medium, and cells were transfected with 3 μg pMD2.G (Addgene, #12259), 5 μg psPAX2 (Addgene, #12260), and 8 μg lentiviral transfer plasmid, diluted in 500 μl Opti-MEM (Thermo Fisher, #11058021), using 48 μl TransIT-Lenti Transfection Reagent (Mirus Bio, #10412), according to the manufacturer’s protocol. Viral supernatants were harvested 60–72 h post-transfection, filtered through a 0.22-μm filter, and concentrated by ultracentrifugation at 61,000 x g for 2 h at 4°C. Viral pellets were resuspended in 100 μl sterile PBS by gentle pipetting. Concentrated lentivirus was used immediately or aliquoted and stored at −80°C.

### Generation of PHF13 knockdown TS cells or knockout TS and BeWo cells

TS cells expressing PHF13 shRNA or control shRNA were generated by lentiviral transduction following our published protocols^30, 69^. Briefly, the PHF13 shRNA lentiviral plasmid (Sigma, #TRCN0000236017), targeting the sequence 5’-TGTCACCTGATCAGGTCAAAG-3’, and a non-targeting control shRNA plasmid (Sigma, #SHC204) were packaged into lentiviral particles and used to transduce TS cells in the presence of polybrene (8 μg/ml). Following transduction, cells were selected with puromycin (5 μg/ml) for 7 days to generate stable knockdown lines.

Candidate guide RNAs (gRNAs) targeting PHF13 were designed as previously described^70, 71^, using the CRISPR/Cas9 or Cas12a technologies. Specifically, TS cells expressing doxycycline-inducible Cas9 and selected with puromycin were generated as previously reported^69^, using pCW-Cas9 (Addgene, #50661)^72^. For Cas9-mediated knockout, the gRNA sequence used (5’-ACTCATTACACTCGATCATG-3’) was cloned into a derivative of the plasmid (Addgene, #57825) modified to contain the blasticidin S deaminase resistance cassette. Following transduction, TS/Cas9 cells expressing the PHF13 gRNA were selected with 5 μg/mL blasticidin. For Cas12a-mediated knockout, the two gRNA sequences were used: gRNA1 (5’-CCGGCCGCCCCATGATCGAGTGT-3’) and gRNA2 (5’-ACATCCGCCGTTCCAACCGCTCG-3’). Oligo gRNAs were cloned into lentiviral backbone (Addgene, #84739) encoding the Cas12a nuclease and a puromycin resistance cassette. Recombinant plasmids were verified by Sanger sequencing. TS or BeWo cells were transduced with lentiviruses and further selected with 5 μg/mL puromycin for seven days to generate stable cell lines.

Knockout clones were generated and screened following the published protocols^30, 69, 70^. Briefly, single cells were sorted into ten 96-well plates to establish clonal populations. Genomic DNA was extracted using QuickExtract DNA solution (Lucigen, #QE09050) according to the manufacturer’s instructions. Target loci were amplified using Q5 hot start high-fidelity 2X master mix (New England Biolabs, #M0494S). Derived PCR amplicons were denatured at 95°C for 10 min and re-annealed to facilitate heteroduplex formation using the following ramp conditions: 95-85°C (ramp rate of -2°C /sec); 85-25°C (ramp rate of -0.3°C /sec). DNA heteroduplex strands were then digested with T7 endonuclease I (New England Biolabs, #M0302S) at 37°C for 1 h to detect insertion/deletion mutations. Clones showing cleavage products were further expanded and validated by immunoblotting to confirm loss of PHF13 protein expression.

### Knockdown of THAP11 in TS cells

Candidate Cas13 gRNA targeting the THAP11 coding region were designed with Cas13design, and the top three guides (gRNA1-3) were selected based on predicted score/accessibility. The sequences of gRNAs are: gRNA1 (5’-AAACGTGTAGAAGTGCAGCGCCT-3’), gRNA2 (5’-ATCTTCACTTCCATCAGAGCCAG-3’), and gRNA3 (5’-ACGCGACACGTTCTTGAGCCAGA-3’). gRNAs were cloned into the pSLQ4419 pHR lentiviral backbone (Addgene, #214875) using BsmBI digestion and NEBuilder HiFi DNA Assembly Master Mix (New England Biolabs, #E2621S). Recombinant plasmids were sequence-verified. For functional validation, gRNA plasmids or a non-targeting control pSLQ7501 (Addgene, #214876) were co-transfected with RfxCas13d pSLQ5428 (Addgene, #155305) into HEK293T/17 cells and knockdown was assessed by qPCR. When compared to a non-targeting control, gRNA1 and gRNA2 showed the best efficiency (50∼60%) and were used for subsequent experiments.

TS cells were transduced with THAP11 gRNA1/gRNA2 or non-targeting control lentivirus in the presence of polybrene, followed by 5 μg/mL puromycin selection for five days to generate gRNA-expressing cells. Cells were then transduced with RfxCas13d lentivirus and cultured for four additional days. THAP11 knockdown and non-targeting control TS cells were reseeded into three separate plates for downstream analyses: (1) a 6-well plate for Western blot analysis to assess THAP11 knockdown efficiency, using an anti-THAP11 antibody (Bethyl Laboratories, #A303-180A, 1:5,000), with histone H3 serving as the loading control (Cell Signaling Technology, #4499S, 1:10,000); (2) a 24-well plate for RNA extraction from TS-CTB cells; and (3) a second 24-well plate for RNA extraction from TS-STB cells, following 48 h of differentiation induction. Once THAP11 knockdown was confirmed by Western blot, total RNA was extracted and shipped to Novogene for RNA library construction and sequencing.

### RNA extraction, library preparation, and RNAseq analyses

Total RNA was extracted from cells using TRIzol RNA isolation reagent (Thermo Fisher, #15596026), followed by purification with the RNeasy mini kit (Qiagen, #74104) including on-column RNase-free DNase (Qiagen, #79254), according to the manufacturer’s protocols as previously described^26^. RNA integrity and quality was assessed using an Agilent HS Total RNA 15-nt kit (Agilent, DNF-472T33) on an Agilent Analytical 5300 Fragment Analyzer. Samples with RNA integrity number (RIN) ≥7 were used for downstream analyses.

RNAseq libraries were prepared by Novogene using poly (A) selection and sequenced on an Illumina NovaSeq 6000 platform to generate paired-end 150-bp reads, with an average depth of ∼40 million reads per sample. After quality control, clean reads were aligned to the human reference genome (GRCh38) using STAR^73^ with default parameters. Uniquely mapped sequencing reads were assigned to GENCODE v47^74^ genes using featureCounts^75^ with the parameters “–p –Q 10 –O”. Genes with read counts more than 10 in at least three of the samples were retained for downstream analysis. Count data were normalized using the TMM (trimmed mean of M values) method and differential gene analysis was performed using edgeR (v3.20.8)^76, 77^. P values were adjusted using the Benjamini-Hochberg procedure^78^ to control the false discovery rate (FDR), and genes with an adjusted p value < 0.05 were considered significantly differentially expressed.

To assess concordance between *PHF13* knockdown datasets (kd1 and kd2), genes were defined as upregulated or downregulated relative to control using an FDR threshold of < 0.05. For each direction (upregulated and downregulated separately), odds ratios were calculated from 2 x 2 contingency tables to quantify enrichment of genes regulated in the same direction in both datasets. An odds ratio greater than 1 indicated that genes significantly upregulated in *PHF13* kd1 were more likely to be upregulated in *PHF13* kd2, suggesting concordant transcriptional effects. Same analysis was performed for downregulated genes. Statistical significance of overlap was evaluated using Fisher’s exact test.

Log_2_ fold changes in gene expression between knockdown and control samples were used for gene set enrichment analysis (GSEA, v4.3.2)^79^. Gene Ontology (GO) and Kyoto Encyclopedia of Genes and Genomes (KEGG) pathway enrichment analyses were performed on differentially expressed genes using DAVID^80, 81^ with an FDR cutoff of 0.05. All expressed genes detected in the RNAseq dataset were used as the background gene se *Cleavage Under Targets & Release Using Nuclease and sequencing (CUT&RUNseq)* CUT&RUN was performed using the CUTANA™ CUT&RUN kit (EpiCypher, #14-1048) according to the manufacturer’s protocol. For each reaction, 0.5 µg of antibodies against PHF13 (Sino Biological, #202944-T08), H3K4me3 (EpiCypher, #13-0060), and a control antibody for

CUT&RUN (EpiCypher, #13-0042) were used. Two biological replicates were performed for each condition. The immunoprecipitated genomic DNA was purified and used for library preparation using the NEBNext Ultra II DNA library prep kits (New England Biolabs, #E7645S). Libraries were sequenced on an Illumina NovaSeq 6000 platform to generate paired-end 150 bp reads, yielding approximately 5 million raw reads per library.

Raw sequencing reads were first processed to remove adapter sequences and low-quality bases. Reads passing quality filtering were aligned to the human reference genome (GRCh38) using the Bowtie 2 (v2.5.2) alignment algorithm in end-to-end mode with default settings^82^. Only uniquely mapped reads were retained using ngsutilsj (v0.5.0), an updated java port of the NGSUtils toolkit^83^, and PCR duplicates were removed using Picard MarkDuplicates (v2.8.16; Picard Toolkit, Broad Institute). The resulting deduplicated BAM files were used for downstream peak calling.

PHF13-enriched regions were identified relative to control IgG libraries using the SICER2 (v1.0.2)^84, 85^. Peaks were called with a window size of 200 bp and a gap size of 600 bp. Significantly enriched PHF13-bound genomic regions were defined using a Bonferroni-adjusted cutoff of p < 0.01. Biological replicates were assessed for concordance based on the overlap of called peaks, and reproducible peaks were used for subsequent target gene annotation.

PHF13 target genes were assigned when a PHF13 peak was located between 10 kb upstream and 500 bp downstream of the transcription start site (TSS) based on GENCODE v38 gene annotation. Genome browser tracks were generated by bedGraphToBigWig (v2.8) and visualized in the UCSC Genome Browser (hg38 assembly). Tracks for individual genes were further visualized using customized R scripts.

Integrated analysis of PHF13 target genes and differentially expressed genes identified in *PHF13* kd TS cells was performed using an over-representation analysis (ORA) algorithms^47^. ORA evaluates whether predefined biological categories such gene subsets and pathways are represented more frequently in a selected gene set than would be expected by chance when compared with a background gene set, e.g all genes expressed in the genome. Fold enrichment quantifies the magnitude of enrichment, representing how much frequently a given biological category appears in the analyzed gene list relative to random expectation. Statistical significance is assessed using a p value, which reflects the probability of observing the same or a greater level of enrichment if the gene list were randomly drawn from the background set.

Motif enrichment analysis of PHF13-bound sequences was conducted using Hypergeometric Optimization of Motif EnRichment (HOMER, v5.1) with findMotifsGenome.pl program against the hg38 genomic background^86^.

### Reverse Transcription and Real-time quantitative PCR (RT-qPCR)

Total RNA was reverse transcribed using High-Capacity cDNA reverse transcription kit (Thermo Fisher, #4368814) according to the manufacturer’s protocols. Quantitative PCR was performed using SYBR select master mix on a QuantStudio 5 Real-Time PCR System (Thermo Fisher).

Relative gene expression changes were calculated using the 2-∆∆CT method^87^ and normalized to the housekeeping genes tyrosine 3-monooxygenase/tryptophan 5-monooxygenase activation protein zeta (*YWHAZ*) and glyceraldehyde-3-phosphate dehydrogenase (*GAPDH*). No-template (H_2_O) controls were included in each experiment. Amplification specificity was confirmed by dissociation curves of the PCR products. Oligo sequences were validated by BLAST (Basic Local Alignment Search Tool for specificity) and provided in Supplementary Table 9.

### Western Immunoblotting

The cells were lysed on ice in RIPA buffer (Millipore, #20-188) supplemented with protease inhibitor cocktail mini EDTA-free and PhoSTOP (Thermo Fisher, #A32961). The protein concentration was quantified by Pierce BCA Protein Assay Kit (Thermo Fisher, #23227). Protein samples (20 µg) were separated on sodium dodecyl sulfate-polyacrylamide gel electrophoresis and transferred onto 0.2 µm polyvinylidene difluoride membrane using standard procedures.

Membranes were immunoblotted with PHF13 (Sino Biological, #202944-T08, 1:2,000), CDX2 (Thermo Fisher, #MA5-35215, 1:6,000), PENK (Thermo Fisher, #MA5-37792, 1:6,000), THAP11 (Bethyl Laboratories, #A303-180A; 1:5,000), histone H3 (Cell Signaling Technology, #4499S; 1:10,000), a mouse monoclonal anti-actin antibody (Sigma, #MAB1501; 1:40,000). A horseradish peroxidase-conjugated goat anti-mouse IgG (Jackson ImmunoResearch, #115035146, 1:40,000) or goat anti-rabbit IgG (Cell Signaling Technology, #7074, 1:10,000) was used as a secondary antibody. The blots were processed for chemiluminescence using the WesternBright Sirius kit (Advansta, #K-12043-D20) and imaged with Chemidoc system.

### hCG ELISA

The hCG ELISA kit (DRG International, #EIA1911R) was used to analyze the hCG levels released into the medium, as we have published^88, 89^. In brief, BeWo cells were cultured in the F-12K complete medium with exposure to the vehicle control or 50 μM forskolin to induce cell fusion. Supernatants of cultured cell medium were collected, and an enzyme immunoassay was performed using the hCG standards to compare hCG proteins in *PHF13* ko *vs* wt cells.

### Statistical analysis and reproducibility

Data are presented as means ± SD where relevant, derived from at least three independent experiments, as indicated in each figure legend. Statistical analyses of RT-qPCR data were performed using Prism, version 10.0 (GraphPad). For two group comparisons, Student’s t-test was used. For all pairwise comparisons, one way ANOVA with *post hoc* Tukey’s test was conducted. Statistical significance was determined as p < 0.05.

## Availability of data and materials

RNAseq data were deposited in NCBI Sequence Read Archive with BioProject IDs: PRJNA1426890, PRJNA1426893, PRJNA1426896, PRJNA1426897.

## Author contributions

Y.O. and Y.S. conceived the project; Y.O. and Y.S. designed the study; Y.O., L.L., J.M., E.S., K.H., and H.S. performed the experiments; Y.O., S.L., T.C. analyzed the data; Y.O., S.L., L.L., and Y.S. reviewed the data and interpreted the results. Y.O. and Y.S. wrote the manuscript. All the authors reviewed the manuscript.

## Acknowledgements

We thank Lori Rideout for assistance in manuscript preparation. We also thank Jose M. Polo for providing us with TS cells.

## Abbreviations

ASCT2: Alanine, serine, cysteine transporter 2, aka SLC1A5
CDX2: Caudal type homeobox 2
CRH: Corticotropin-releasing hormone
CTB: Cytotrophoblast
CUT&RUNseq: Cleavage under targets & release using nuclease and sequencing
CYP11A1: Cytochrome P450 family 11 subfamily A member 1
CYP19A1: Cytochrome P450 family 19 subfamily A member 1 DEG Differentially expressed gene
ELF5: E74 like E26 transformation-specific transcription factor 5
ERVW-1: Endogenous retrovirus family W 1 (aka syncytin-1, ERVWE1)
ERVFRD-1: Endogenous retrovirus family FRD 1 (aka syncytin-2, ERVFRDE1)
GAPDH: Glyceraldehyde-3-phosphate dehydrogenase
GATA2/3: GATA binding protein 2/3
GCM1: Glial cells missing transcription factor 1
H3K4me3: Histone 3 lysine 4 trimethylation
hCG: Human chorionic gonadotropin
HMOX1: Inducible heme oxygenase 1
LGALS13/14/16: Lectin, galactoside-binding, soluble, 13/14/16, aka galectin 13/14/16
MFSD2a: Major facilitator superfamily domain containing 2a
OVOL1: Ovo like transcriptional repressor 1
PGF: Placental growth factor
PHF13: Plant homeodomain finger protein 13
PSG3/11: Pregnancy specific beta-1-glycoprotein 3/11
SDC1: Syndecan 1
SLC1A5: Solute carrier family 1 member 5
SOX15: SRY-box transcription factor 15
STB: Syncytiotrophoblast
TBX3: T-box transcription factor 3
TEAD1/3/4: Transcriptional enhancer factor-1 domain transcription factor 1/3/4
TET1/2: Ten-eleven translocation methylcytosine dioxygenase 1/2 (TET1/2)
TFAP2A/C: Transcription factor activator protein AP-2 alpha/gamma
THAP11: Thanatos-associated protein domain 11
TS: Trophoblast stem
TP63: Tumor protein p63
YAP1: Yes-associated protein 1
YWHAZ: Tyrosine 3-monooxygenase/tryptophan 5-monooxygenase activation zeta

## Supplementary Note 1

Villous trophoblast differentiation is associated with decreased transcripts of *opioid receptors* (*OPRs)*, including *OPRM1, OPRD1,* and *OPRK1*^26^, and the loss of *PHF13* promoted trophoblast differentiation, we therefore examined whether PHF13 regulates OPR pathway-relevant genes, including OPRs and opioid peptide agonists. Although *OPRD1*, *OPRK1*, and *OPRL1* were expressed in TS-CTB, their transcripts were unchanged in *PHF13* kd cells. We noted that *OPRM1* mRNA was not detected by RNAseq, consistent with low abundance based on our recent qPCR analysis^26^. These results suggested that PHF13 does not directly regulate OPR transcription, but prompted us to test whether PHF13 instead controls OPR pathway activity by regulating the precursor genes encoding opioid peptide agonists. These include prodynorphin (PDYN), which generates dynorphins acting primarily on OPRK1; proopiomelanocortin (POMC), which produces β-endorphin preferentially stimulating OPRM1; and proenkephalin (PENK), which generates enkephalin peptides mainly activating OPRD1.

*PDYN* was undetected by RNAseq and *POMC* was not affected by PHF13. In contrast, *PENK* mRNA was strongly reduced in *PHF13* kd cells (log_2_FC: -4.58). Consistent with direct regulation, we found that PHF13 occupied the *PENK* genomic locus in TS-CTB, but not in TS-STB. Nevertheless, PENK protein levels in TS-STB were comparable to TS-CTB (Suppl. Fig. 5). Together, these results imply that PHF13 is required to sustain PENK expression in TS-CTB, with regulation in TS-STB likely maintained by PHF13-independent mechanisms. This state-specific regulation of PENK exemplifies the broader principle that PHF13 engages distinct genomic programs depending on trophoblastic cell states.

## LEGENDS FOR SUPPLEMENTAL FIGURES

**Suppl. Fig. 1:**
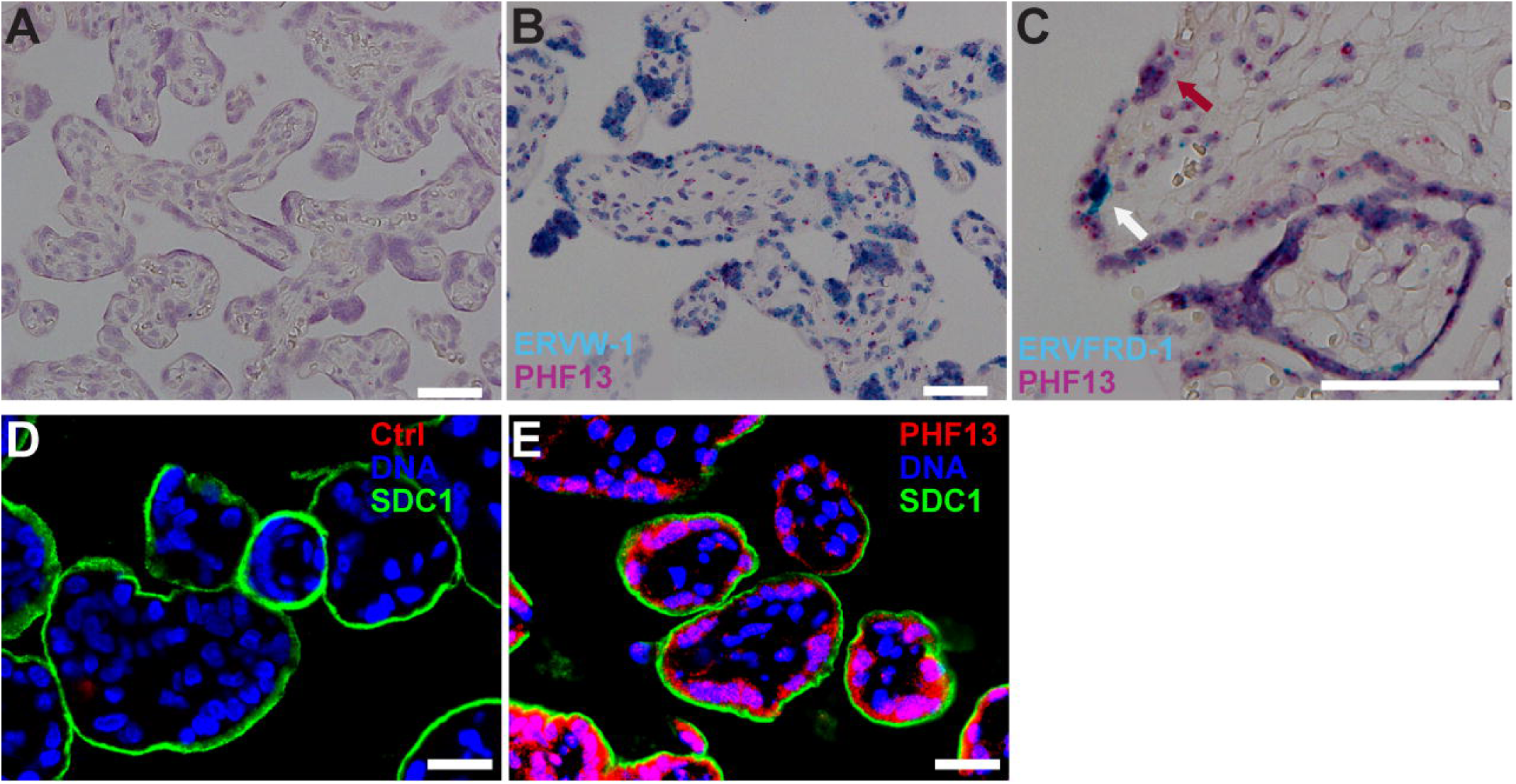
The expression of PHF13 in healthy term placental specimens. **A-C.** RNAscope mRNA analysis was peformed using validated RNA probes, including negative control probe (A), *ERVW-1* (B), *ERVFRD-1* (C) and *PHF13* (B-C). **D-E.** PHF13 protein, analyzed using the multiplexed ChipCytometry, including controls (D) and PHF13 (E). Note that the syncytiotrophoblast layer was marked with SDC1, and nuclei were counterstained with the Hoechst 33342 DNA dye. Scale bars: 10 μm. Images present 3 independent experiments.

**Suppl. Fig. 2:**
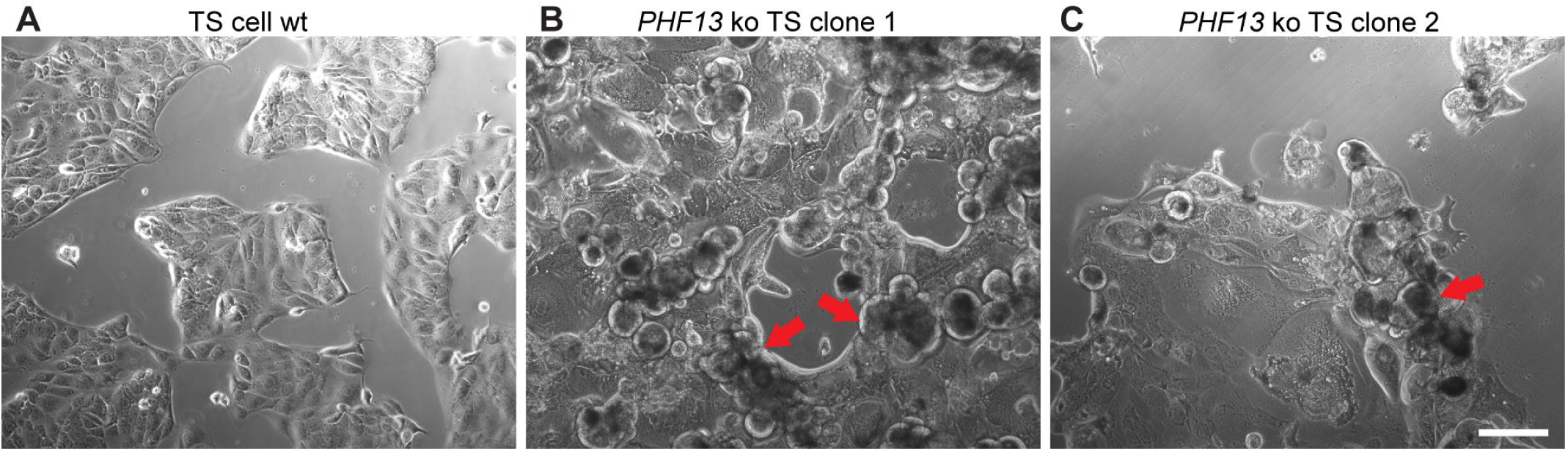
Loss of PHF13 in trophoblast stem cells reduces cell viability. Wild-type (wt) and PHF13 knockout (ko) cells were visualized by the bright-field microscopy. Compared to control cells (A), which show the expected cobblestone-like morphology, two ko clonal cells (B-C) exhibit overt cell death (red arrows). Scale bars: 10 μm.

**Suppl. Fig. 3:**
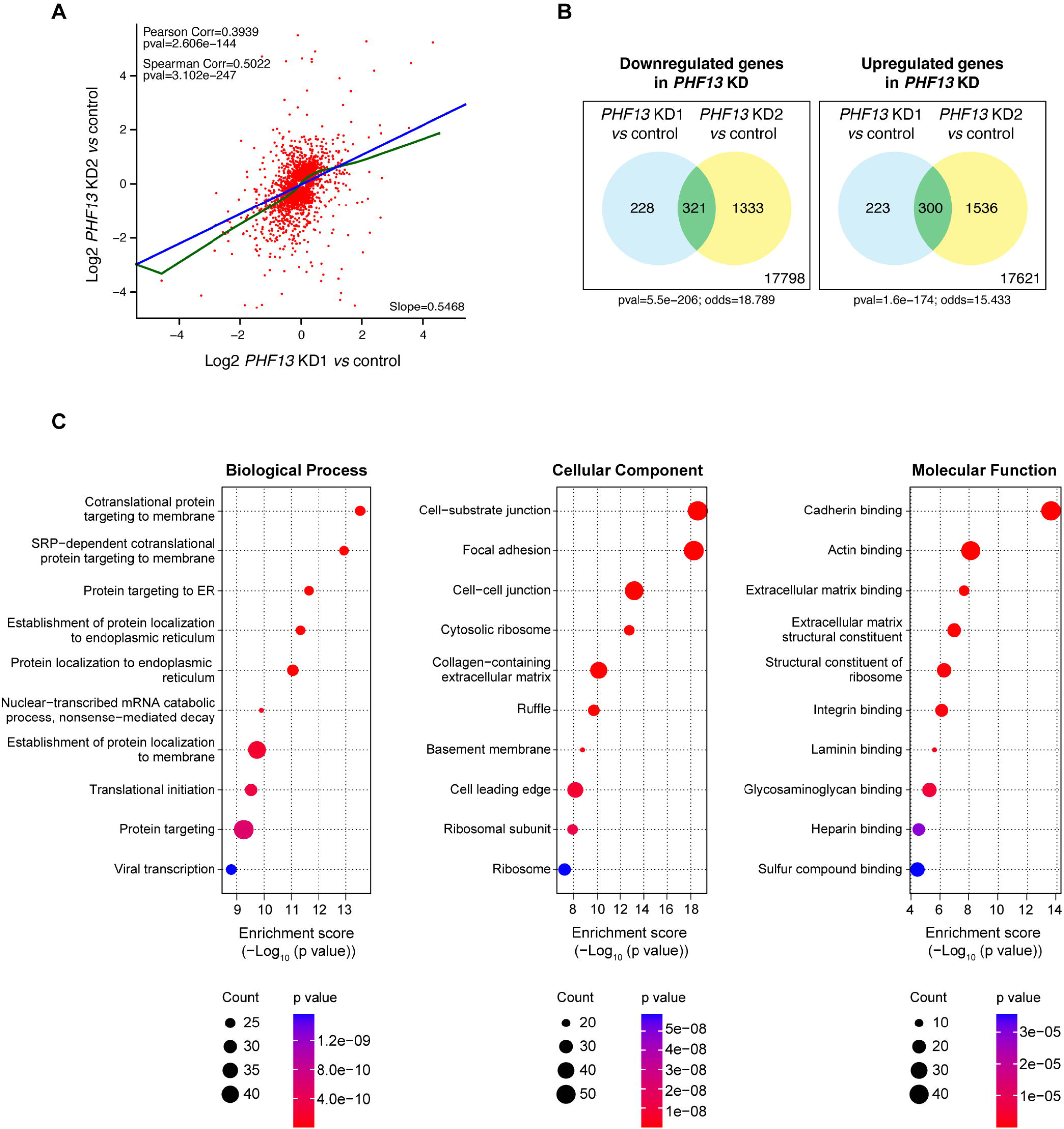
Correlation and gene ontology analyses of transcriptomic changes in two independent PHF13 knockdown TS cell lines. **A.** Correlation of gene expression changes between *PHF13* kd1 and *PHF13* kd2 TS cells relative to their respective controls. The X-axis represents the log_2_ fold change in gene expression (*PHF13* kd1 vs control), whereas the Y-axis shows the log_2_ fold change in gene expression (*PHF13* kd2 vs control). Significant p values define both the Pearson and Spearman correlation analyses. The blue line indicates linear regression, whereas the green line shows non-linear regression. **B.** Venn diagrams showing overlap of differentially expressed genes in two PHF13 kd lines, when compared to their controls. 321 downregulated and 300 upregulated genes were found in both kd lines. The odd ratios for overlap were 18.79 for the downregulated genes, and 15.43 for the upregulated genes, suggesting signficant concordance between the two knockdown lines. ***C.*** Gene ontology (GO) analysis of the 621 concordantly regulated genes shared in both PHF13 kd TS cells. The top 10 enriched terms are shown for each GO category, including biological processes, celluar component, and molecular function, ranked by enrichment scores. See results for details.

**Suppl. Fig. 4:**
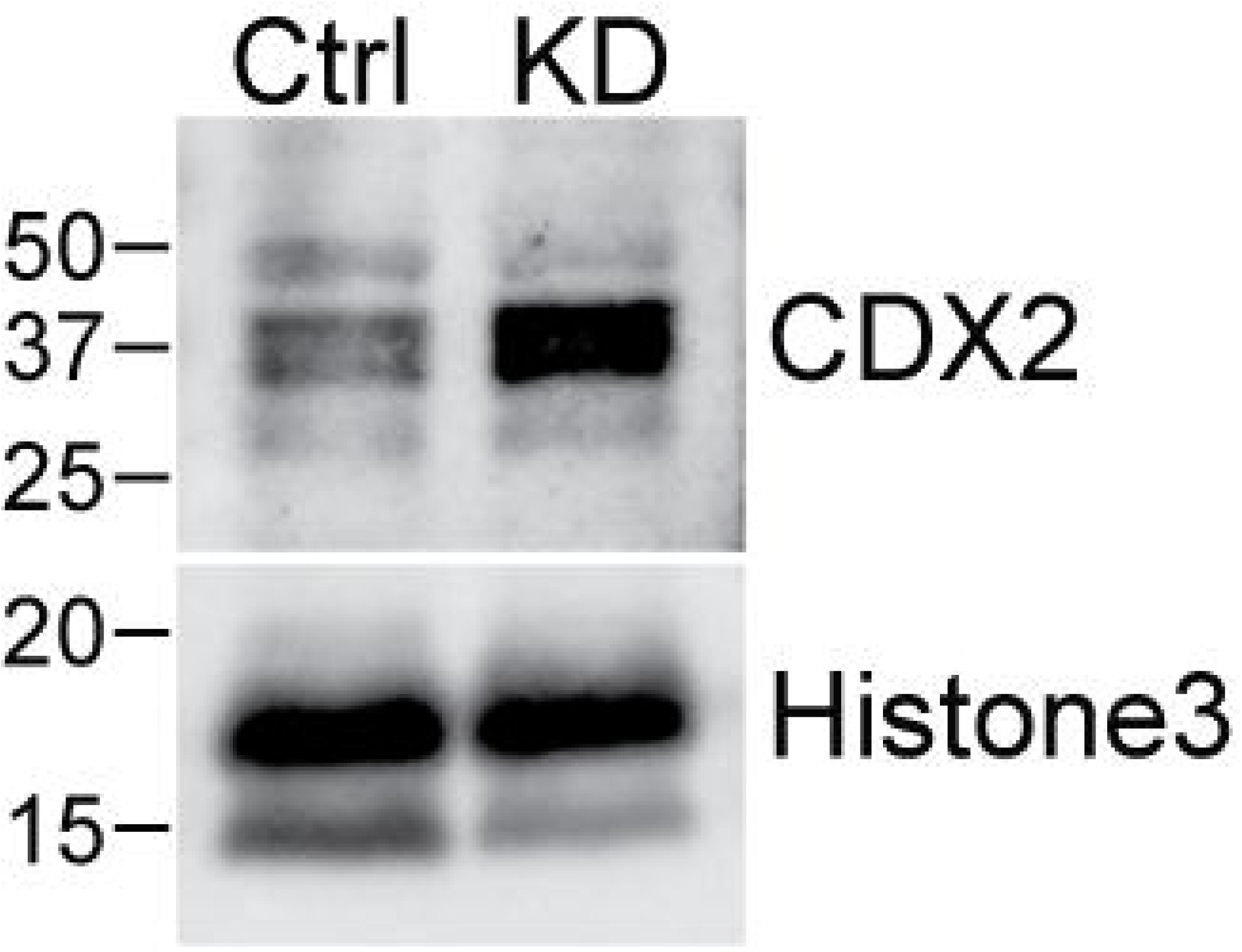
CDX2 protein is elevated in PHF13 knockdown TS-CTB cells. CDX2 levels were assessed by immunoblotting using a rabbit anti-CDX2 antibody, and histone H3 was used as a loading control (n=3).

**Suppl. Fig. 5:**
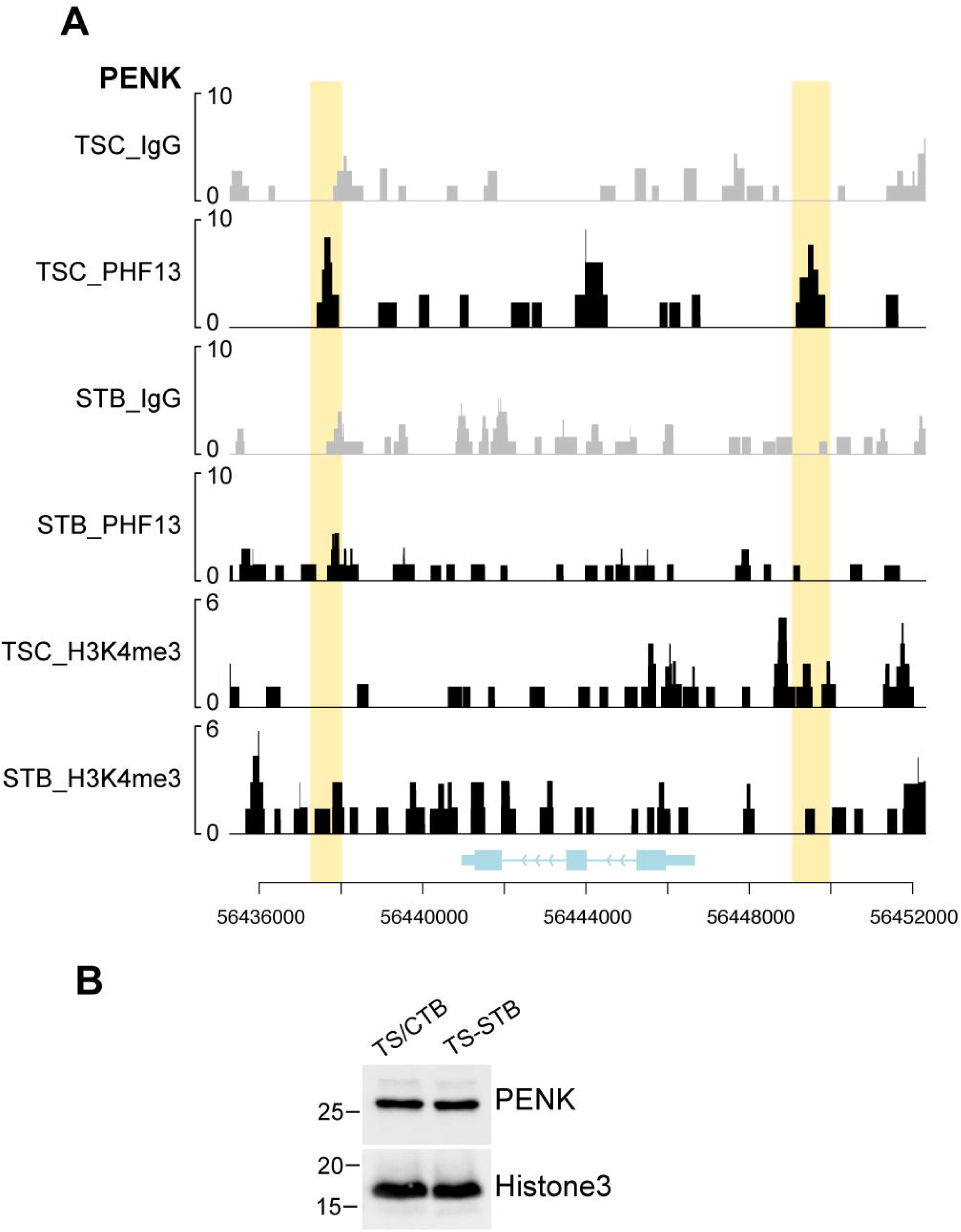
PHF13 promotes PENK expression in TS-CTB cells. **A.** PHF13 binds the *PENK* locus in TS-CTB but not in TS-STB. Yellow shading indicates significant PHF13 peaks relative to IgG controls; no H3K4me3 peaks were detected in these shaded regions in TS-CTB. The x-axis represents genomic corordinates with exonic and intronic structure, and arrows indicate transcription direction. The y-axis represents normalized peak intensity. **B.** PENK protein is expressed in both TS-CTB and TS-STB, and histone H3 was used as a loading control (n=3).

